# Analysis of continuous neuronal activity evoked by natural speech with computational corpus linguistics methods

**DOI:** 10.1101/2020.04.21.052720

**Authors:** Achim Schilling, Rosario Tomasello, Malte R. Henningsen-Schomers, Alexandra Zankl, Kishore Surendra, Martin Haller, Valerie Karl, Peter Uhrig, Andreas Maier, Patrick Krauss

## Abstract

In the field of neurobiology of language, neuroimaging studies are generally based on stimulation paradigms consisting of at least two different conditions. Designing those paradigms can be very time-consuming and this traditional approach is necessarily data-limited. In contrast, in computational linguistics analyses are often based on large text corpora, which allow a vast variety of hypotheses to be tested by repeatedly re-evaluating the data set. Furthermore, text corpora also allow exploratory data analysis in order to generate new hypotheses. By drawing on the advantages of both fields, neuroimaging and corpus linguistics, we here present a unified approach combining continuous natural speech and MEG to generate a corpus of speech-evoked neuronal activity.

## Introduction

Contemporary linguistic research is characterized by a great variety of methodological approaches. The fields of psycholinguistics and neurobiology of language are characterized by a vast number of different methods that are applied in order to investigate the neuronal and cognitive correlates and processing principles of language acquisition, representation, comprehension and production [1]. Besides functional magnetic resonance imaging (fMRI) studies [2–4], electrophysiological measurements, i.e. magnetoencephalography (MEG) [5] and electroencephalography (EEG) [6,7], are widely used in neurolinguistics to investigate the neural correlates underlying language processing in the human brain [8–13].

However, most of the experimental studies on language processing conducted so far have focused on one aspect of linguistic information at a time. For instance, neurocognitive studies have explored the neural responses of words compared to pseudo words [14, 15], between different conceptual semantic categories [16], complex against simple grammatical sentences [17], or during pragmatic processing of different communicative actions [13]. Although, all these studies shed light on the correlates of language processing in the human brain, it is still not fully understood whether similar brain responses during single words or sentence understanding also emerge during perception of natural speech, similar to everyday experience. However, recently a growing number of approaches address this issue [4, 18–21].

Furthermore, traditional experimental designs typically consist of at least two different conditions studied under carefully controlled circumstances [10–12]. The measured data are then pre-processed, i.e. referenced, filtered, epoched and averaged, and finally contrasted according to the different stimulation conditions [1]. To obtain a good signal to noise ratio (SNR) of the acquired brain responses, each of these conditions must contain dozens of different items or stimulus repetitions.

For instance, the evaluation of event related potentials (ERPs) from the EEG data, or, in the case of MEG, event related fields (ERFs), requires a relatively large number of stimuli (40-120 trials) per condition to achieve high SNR and ensure sufficient statistical power. This is due to the fact that the signal of a specific condition remains constant across multiple repetitions while the noise signal which is assumed to be randomly distributed, is reduced when large number of time-locked stimuli are pooled together [22–26].

However, creating a large number of stimuli to increase SNR is associated with a serious drawback. It is well known that repeated presentation of a stimulus causes a diminished neural activation, a phenomenon for which the term *repetition suppression* has been coined [27–31]. In fMRI, repetition suppression is observed as a reduced blood oxygen-level-dependent (BOLD) response elicited by a repeated stimulus, also called fMRI adaptation [32]; for a recent review see also [33]. The underlying neuronal mechanisms are still a matter of debate, and range from neuronal fatigue [28], or neuronal sharpening [34], through neuronal facilitation [28] as relatively automatic bottom-up mechanisms, to predictive coding [35]. There, top-down backward influences from higher to lower cortical layers modulate processing in case of a correct prediction of the upcoming stimulus. Hence, repetition suppression reflects a smaller prediction error for expected stimuli, i.e. decreased activation for repeated stimuli. Thus, in order to prevent repetition suppression, it is necessary to design a certain number of different stimuli from each condition to avoid repetition, which is often very challenging or even impossible. One strategy is to focus on single-item ERPs/ERFs, but in such cases it is necessary to compensate by testing more participants to obtain stable signals [36].

Here, we present an alternative approach to overcome the aforementioned limitations of the electrophysiological assessment of language processing, and to open up the possibility of investigating different levels of linguistic information during natural speech comprehension within a single experiment. In particular, in the present study, we investigated brain responses elicited during listening to the audio book edition of a German-language novel by means of MEG measurements (for similar approaches see [3, 37]). Repetition suppression is not expected to occur here, as the same linguistic utterance is not repeatedly presented among a few stimuli types, and if repetition happens, it does in different linguistic contexts (i.e., it possibly occurs with different linguistic units) and also more sparsely.

Other previously published papers describe the use of continuously written stimuli in reading studies while recording EEG/MEG [38–41]. Remarkable, it turned out that the representation of semantic information across human cerebral cortex during listening versus reading is invariant to stimulus modality [4]. Since listening to an audio book during the one-hour measurement session seems to be less strenuous for the participants than reading for the same period of time, we chose acoustic stimulation rather than visual stimulation.

Using computational corpus linguistics (CCL) [42, 43] applied to the analyses of large text corpora, which usually consist of hundreds of thousands or even billions of tokens [44–49], offers the opportunity to test a vast number of hypotheses by repeatedly re-analyzing the data [50] and to deploy modern machine learning techniques on such datasets [51]. Furthermore, text corpora also allow for exploratory data analyses in order to generate new hypotheses [52]. In our approach we can generate a large database of neuronal activity in a single measurement session, corresponding to the comprehension of several thousands of words of continuous speech similarly to everyday language. However, for later studies the data set has to be split into multiple parts (e.g., development/training/test or training/validation/test) in analogy to standard machine learning data sets as MNIST (50,000 training images, 10,000 test images [53]), as hypothesis generation and checking for statistical significance have to be done in two disjoint steps, in order to prevent *HARKing* (hypothesizing after the results are known) [54]. In such cases, inferential statistical analysis is not valid and applicable [55]. Thus, this approach is only possible with a large dataset.

Here, we provide the proof-of-principle of this approach by calculating ERFs and normalized power spectra of word onsets and offsets overall as well as for the group of content words (nouns, verbs, adjectives) and for the group of function words (determiners, prepositions, conjunctions), which are known to differ semantically to a substantial extent. Hence, greater activation for content compared to function words can be expected, as reported in previous studies (e.g., [56, 57])^1^. Furthermore, we check for consistency of the data, by comparing intra-individual differences of neural activity in different brain regions and we perform non-parametric cluster permutation tests to determine significant differences between conditions.

## Methods

### Human participants

Participants were 15 (8 females, 7 males) healthy right-handed (augmented laterality index: *µ* = 85.7, *σ* = 10.4) and monolingual native speakers of German aged 20-42 years. They had normal hearing and did not report any history of neurological illness or drug abuse. They were paid for their participation after signing an informed consent form. Ethical permission for the study was granted by the ethics board of the University Hospital Erlangen (registration no. 161-18 B). For the questionnaire based assessment and analysis of handedness we used the Edinburgh Inventory [58].

### Speech stimuli and natural language text data

As natural language text data, we used the German novel *Gut gegen Nordwind* by Daniel Glattauer (*Q*c *Deuticke im Paul Zsolnay Verlag*, Wien 2006) which was published by *Deuticke Verlag*. As speech stimuli, we used the corresponding audio book which was published by *Hörbuch Hamburg*. Both the novel and the audio book are available in stores, and the respective publishers gave us permission to use them for the present and future scientific studies.

Book and audio book consist of a total number of 40460 tokens (number of words) and 6117 types (number of unique words). The distribution of single word classes and bi-gram word class combinations occurring in the (audio) book were analysed and compared to a number of German reference corpora [59], and in addition, other German novels, by applying *part-of-speech (POS) tagging* [60–62] as implemented in the python library *spaCy* [63]. The similarities, or dissimilarities respectively, of all distributions are visualized using *multi-dimensional scaling (MDS)* [64–67].

The total duration of the audio book is approximately 4.5 hours. For our study, we only used the first 40 minutes of the audio book, divided into 10 parts of approximately 4 minutes (*µ* = 245 *s, σ* = 39 *s*). This corresponds to approximately 6000 words, or 800 sentences, respectively of spoken language, where each sentence consists on average of 7.5 words and has a mean duration of 3 seconds.

In order to avoid cutting the text in the middle of a sentence or even in the middle of a word, we manually cut at paragraph boundaries, which resulted in more meaningful interruptions of the text. For the present study, only the first 3 sections (roughly 12 minutes of continuous speech) of the recordings and the corresponding measurements were analyzed.

### Stimulation protocol

The continuous speech from the audio book was presented in 10 subsequent parts (cf. above) at a sensory level of approximately 30-60 dB SPL. The actual loudness varied from participant to participant. It was chosen individually to ensure good intelligibility during the entire measurement, but also to prevent it from being unpleasant. Simultaneously with auditory stimulation, a fixation cross at the centre of the screen was presented all the time to minimize artifacts from eye movements. After each audio book part, three multiple-choice questions on the content of the previously presented part were presented on the screen in order to test the participants’ attention. Participants had to answer the questions by pressing previously defined keys on a MEG-compatible keyboard. MEG recording was stopped during the question blocks, since these short breaks were also used to allow participants to move and make themselves comfortable again. Furthermore, stimulation was interrupted for a short break of approximately 5 minutes after audio book parts number 4 and 7. The total duration of the protocol is approximately one hour. The complete stimulation protocol is shown in Figure 1.

**Figure 1:**
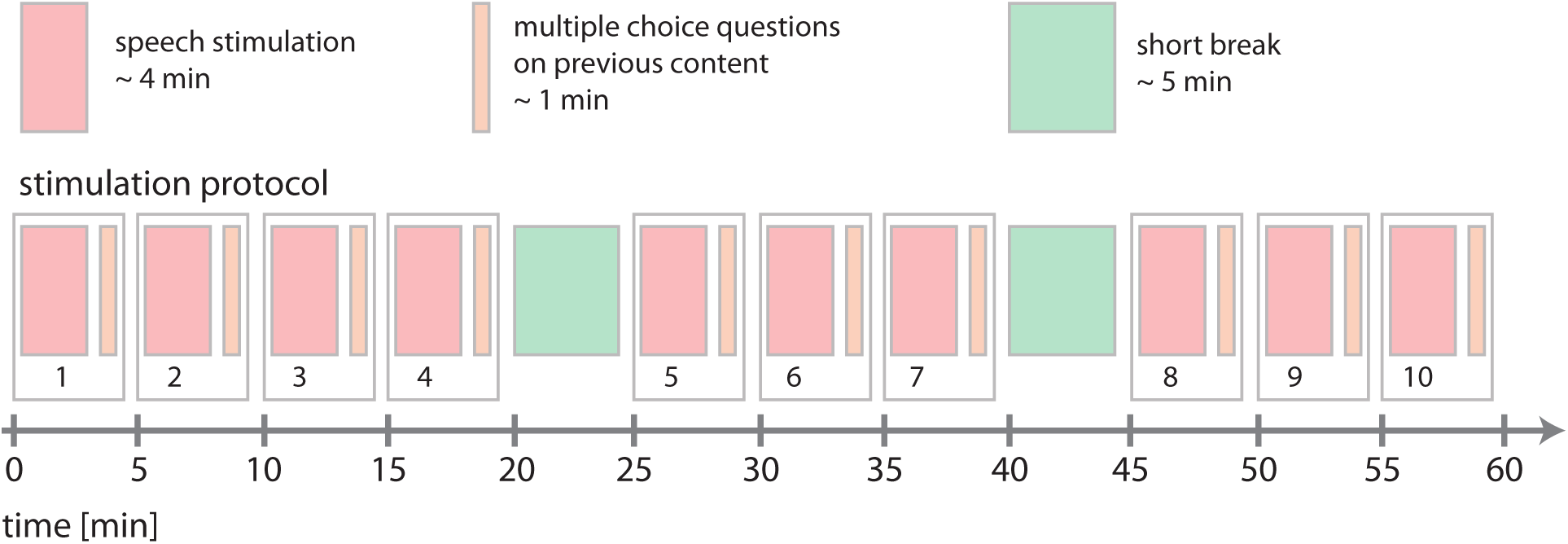
Stimulation protocol. The total duration of the protocol was approximately one hour. The audio book was presented in 10 subsequent parts with an average duration of 4 minutes (red). After each part, three multiple-choice questions on the content of the previous part of the audio book were presented (orange). After audio book parts number 4 and 7, stimulation was interrupted for a short break of approximately 5 minutes (green).

### Generation of trigger pulses with forced alignment

In order to automatically create trigger pulses for both, the synchronization of the speech stream with the MEG recordings, and to mark the boundaries of words, phonemes, and silence for further segmentation of the continuous data streams, *forced alignment* [68–70] was applied to the text and recording. For this study we used the free web service *WebMAUS* [71, 72]. It takes a wave file containing the speech signal, and a corresponding text file as input and gives three files as output: the time tags of word boundaries, a phonetic transcription of the text file, and the time tags of phone boundaries. Even though forced alignment is a fast and reliable method for the automatic phonetic transcription of continuous speech, we carried out random manual inspections in order to ensure that the method actually worked correctly. Although forced alignment is not 100% reliable, manual spot checks found no errors in our alignment. Of course, the high-quality recording of an audio book is among the best possible inputs for such software.

For simplicity, we only used the time tags of word boundaries in this study. However, a more fine grained analysis on the level of speech sounds could easily be performed retrospectively, since the time tags of beginning and ending of a given word correspond to the beginning of the word’s first phone and the ending of the word’s last phone, respectively. Thus, the two lists containing the time tags of the word and phoneme boundaries can easily be aligned with each other.

### Speech presentation and synchronisation with MEG

The speech signal was presented using a custom made setup (Figure 2). It consists of a stimulation computer connected to an external USB sound device (Asus Xonar MKII, 7.1 channels) providing five analog outputs. The first and second analog outputs are connected to an audio amplifier (AIWA, XA-003), where the first output is connected in parallel to an analog input channel of the MEG data logger in order to enable an exact alignment of the presented stimuli and the recorded MEG signals (cf. Figure 2 a). In addition, the third analog output of the sound device is used to feed the trigger pulses derived from forced alignment into the MEG recording system via another analog input channel. In doing so, our setup prevents temporal jittering of the presented signal caused by multi-threading of the stimulation PC’s operating system, for instance. For an overview of the wiring scheme of all devices see Figure 2a.

**Figure 2:**
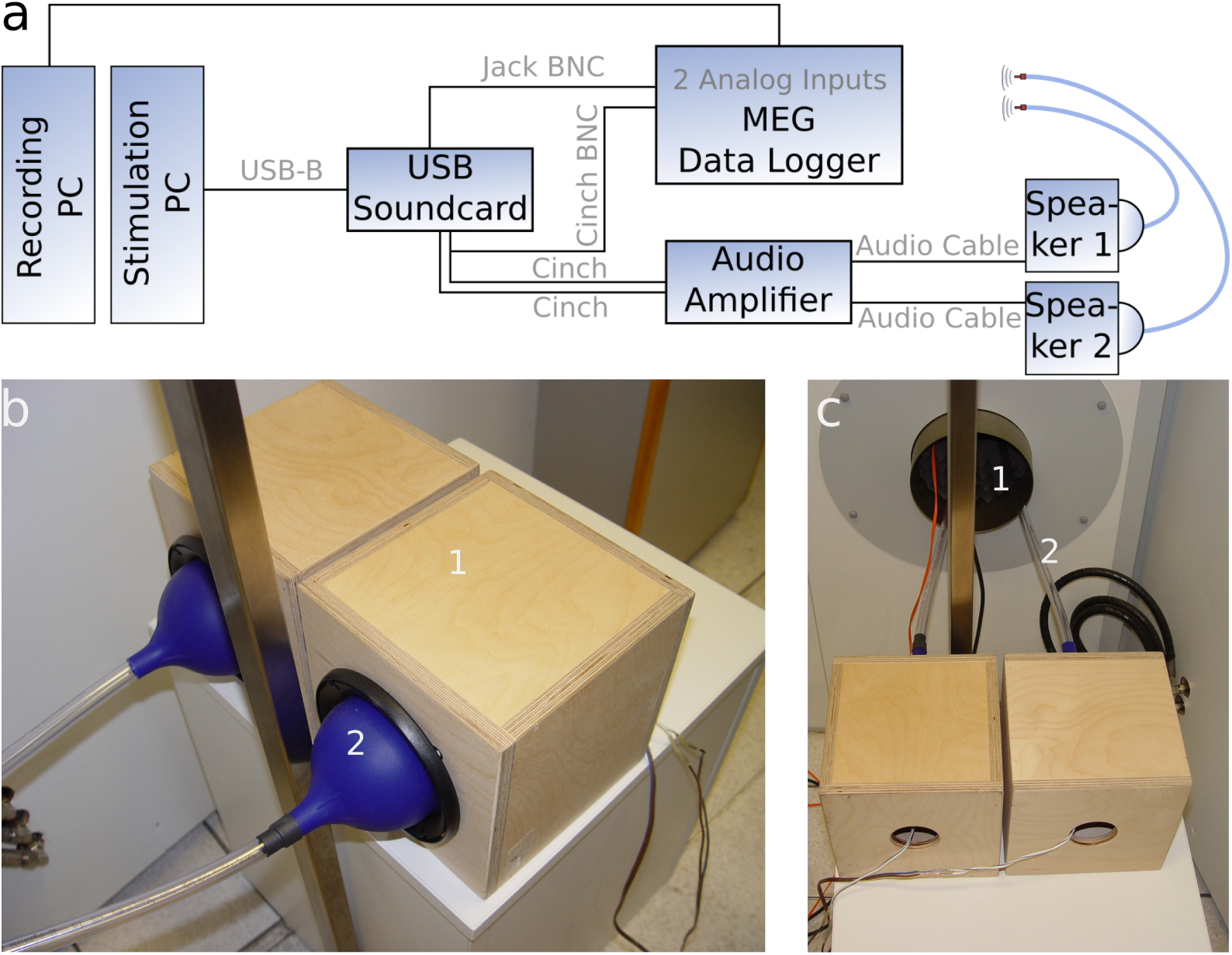
Setup configuration. a: Wiring scheme of the different devices. b: The speech sound is transmitted into the magnetically shielded chamber via a custom-made construction consisting of two loudspeakers (1) which are coupled to silicone funnels (2) each connected to a flexible tube. c: Through a small whole in the magnetically shielded chamber (1), speech sound is transmitted via the two flexible tubes (2).

The speech sound was transmitted into the magnetically shielded MEG chamber to the participants’ ears via a custom-made device consisting of two loudspeakers (Pioneer, TS-G1020F) which are coupled to silicone funnels each connected to a flexible tube of ≈ 2 *m* length and with an inner diameter of ≈ 2 *cm* (Figure 2b). These tubes are led through a small hole in the magnetically shielded chamber to prevent artifacts produced by interfering magnetic fields generated by the loudspeakers (Figure 2c). We carried out calibration tests to ensure that the acoustical distortions caused by the tube system do not affect speech intelligibility. Furthermore, due to the length of the tubes and the speed of sound, there is a constant time delay from the generation of sound to the arrival of the sound at the participant of ≈ 6 *ms*, which we took into account for the alignment described below.

The stimulation software is implemented using the programming language *Python 3.6*, together with Python’s sound device library, the *PsychoPy* library [73, 74] for the stimulation protocol, and the *NumPy* library for basic mathematical and numerical operations.

### Magnetoencephalography and data processing

MEG data (248 magnetometers, 4D Neuroimaging, San Diego, CA, USA) were recorded (1017.25 Hz sampling rate, filtering: 0.1-200 Hz analogue band pass, supine position, eyes open) during speech stimulation. Positions of five landmarks (nasion, LPA, RPA, Cz, inion) were acquired using an integrated digitizer (Polhemus, Colchester, Vermont, Canada). MEG data were corrected for environmental noise using a calibrated linear weighting of 23 reference sensors (manufacturers algorithm, 4D Neuroimaging, San Diego, CA, USA).

Further processing was performed using the Python library *MNE* [75, 76]. Data were digitally filtered offline (1-10 Hz bandpass for ERF analyses; 50 Hz notch on for power spectra analysis) and downsampled to a sampling rate of 1000 Hz. MEG sensor positions were co-registered to the ICBM-152 standard head model [77] and atlas 19-21 [78], as individual MRI data sets for the participants were not available. Furthermore, recordings were corrected for eye blinks and electrocardiography artifacts based on signal space projection of averaged artifact patterns, as implemented in *MNE* [75, 76].

Additionally, we performed an independent component analysis (ICA) and deleted the first two independent components of the data, to further improve data quality. However, it appears that this processing step does not affect the observed differences of neural responses to function and content words (Figures 8 and S8).

**Figure 3:**
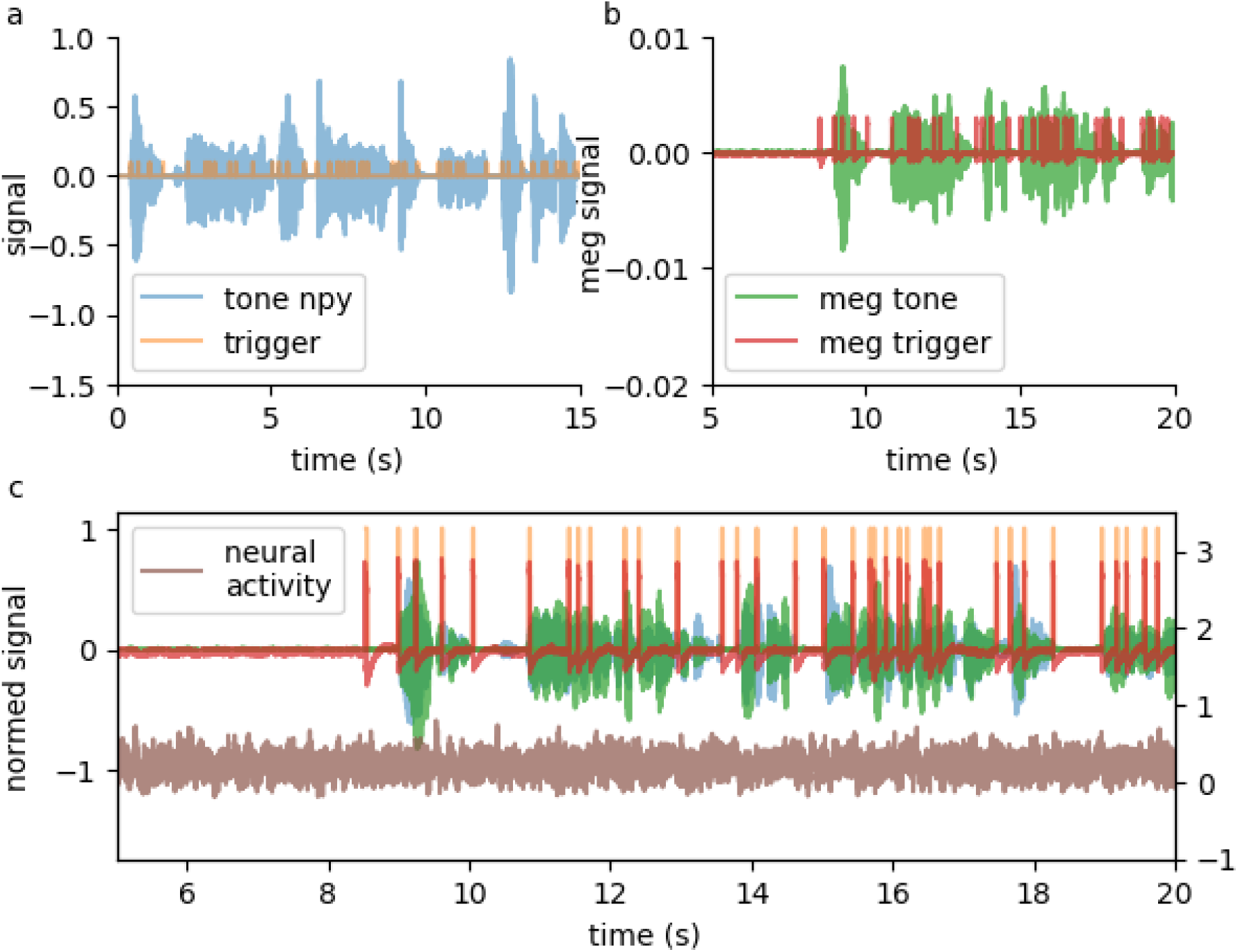
Alignment of speech stream and MEG signal. a: Sample audio book wave file (blue) together with time tags of word boundaries (orange) from forced alignment. b: Corresponding recordings of two analog auxiliary channels of the MEG. c: Alignment of data streams from a and b, together with one sample MEG channel.

**Figure 4:**
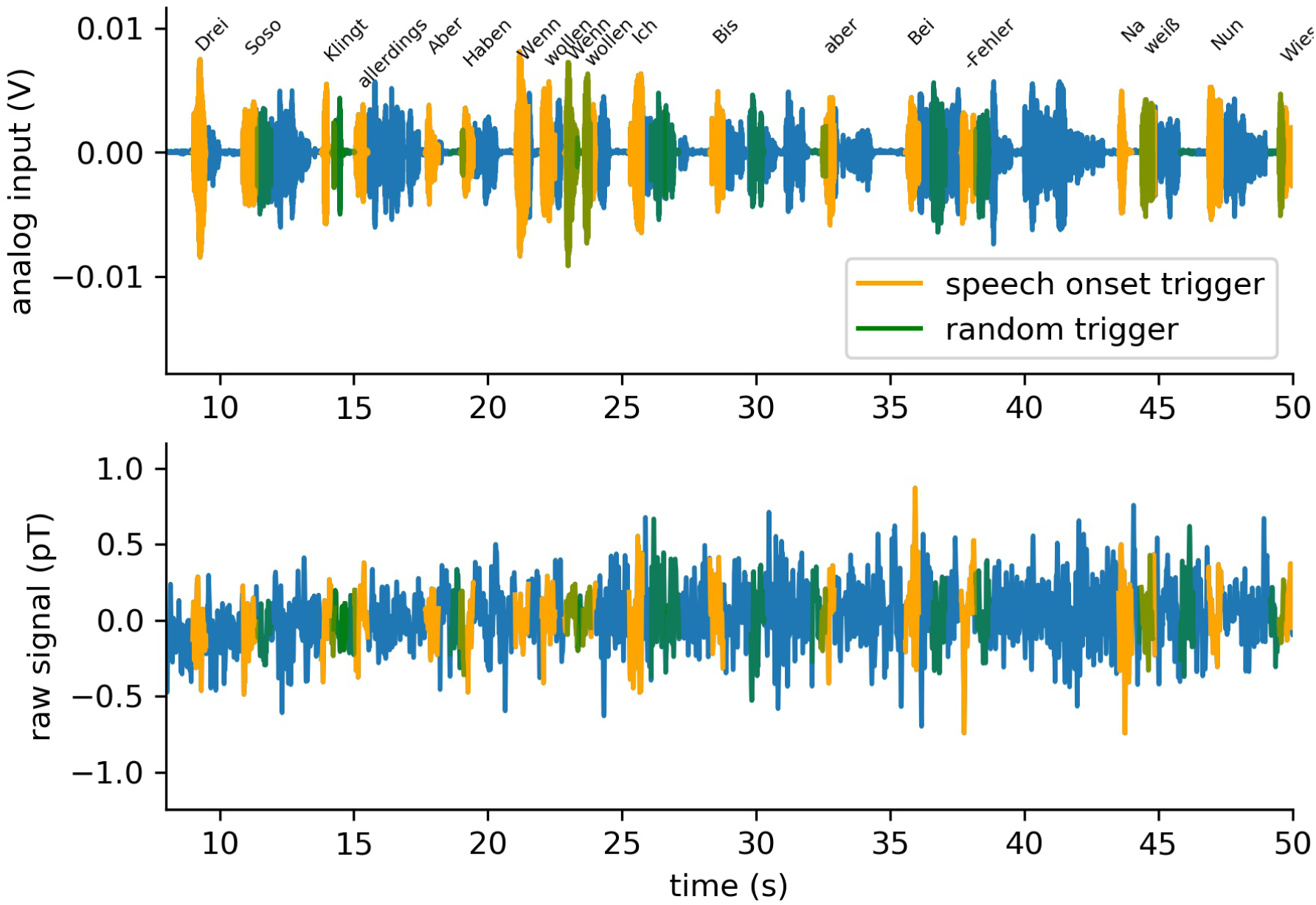
Segmentation of speech stream and MEG signal. After alignment, the continuous wave file (top panel) and multi-channel MEG recordings (bottom panel) are segmented using the time tags from forced alignment as boundaries, and labeled with the corresponding types, i.e. words.

**Figure 5:**
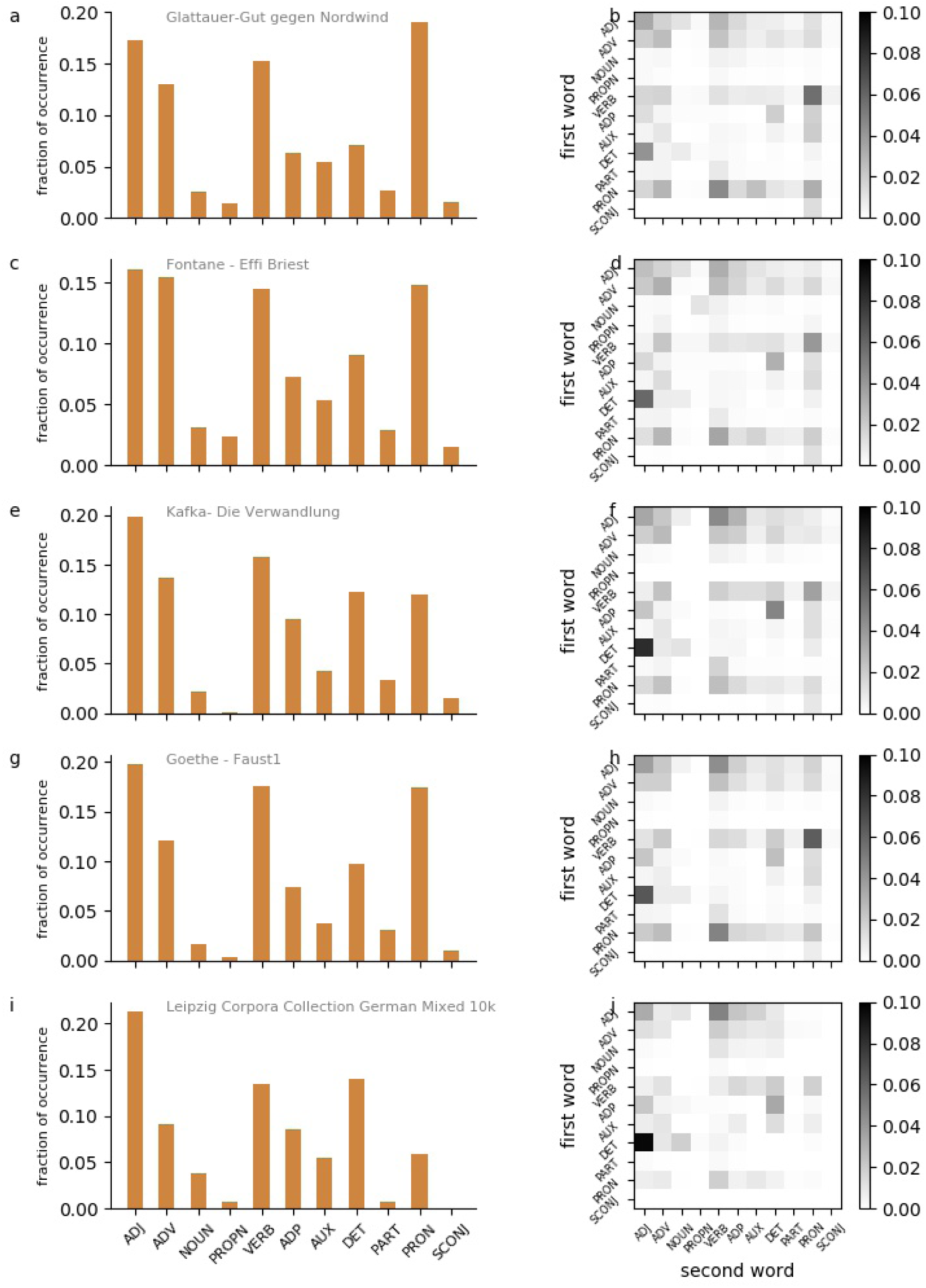
Distributions of word classes and bi-gram word classes. a,c,e,g,i: Distribution of word classes according to POS-tagging. Adjectives (ADJ), adverbs (ADV), nouns (NOUN), proper nouns (PROPN), verbs (VERB), adpositions (ADP), auxiliary verbs (AUX), determiners (DET), particles (PART), pronouns (PRON), subordinating conjunctions (SCONJ). b,d,f,h,j: Distribution of word classes of 2-word sequences. Rows: word class of first word. Columns: word class of second word.

**Figure 6:**
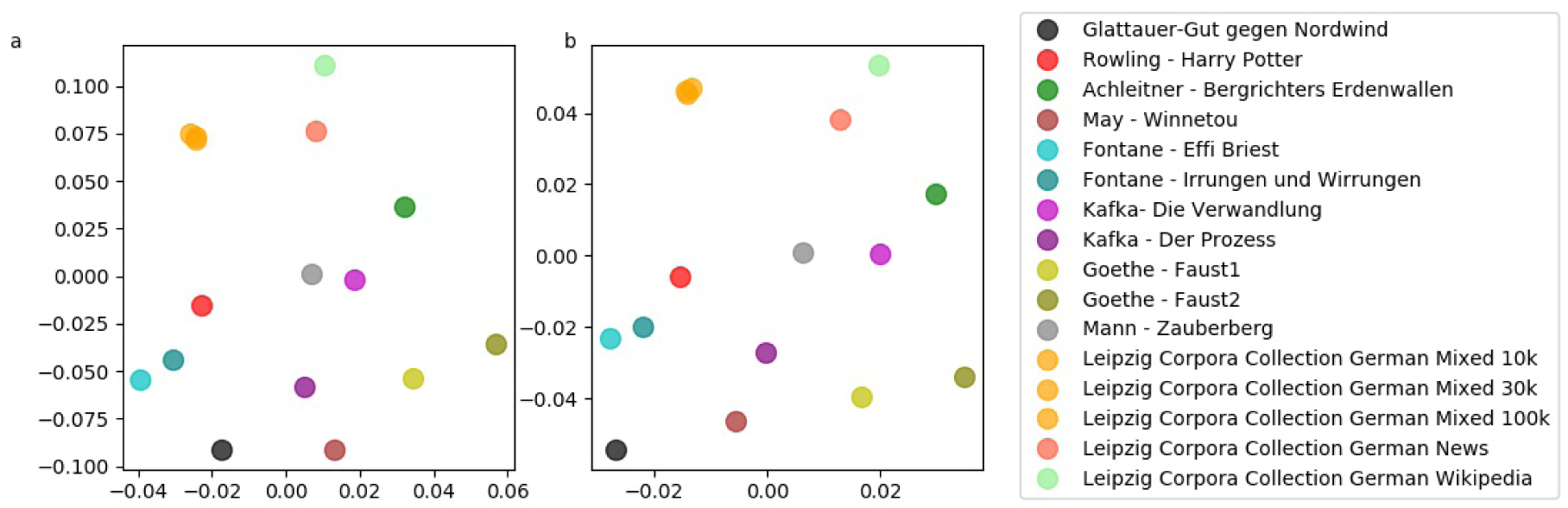
MDS projection of word class distributions. a: MDS projection of distributions of single word classes. b: MDS projection of distributions of bi-gram word classes combinations.

**Figure 7:**
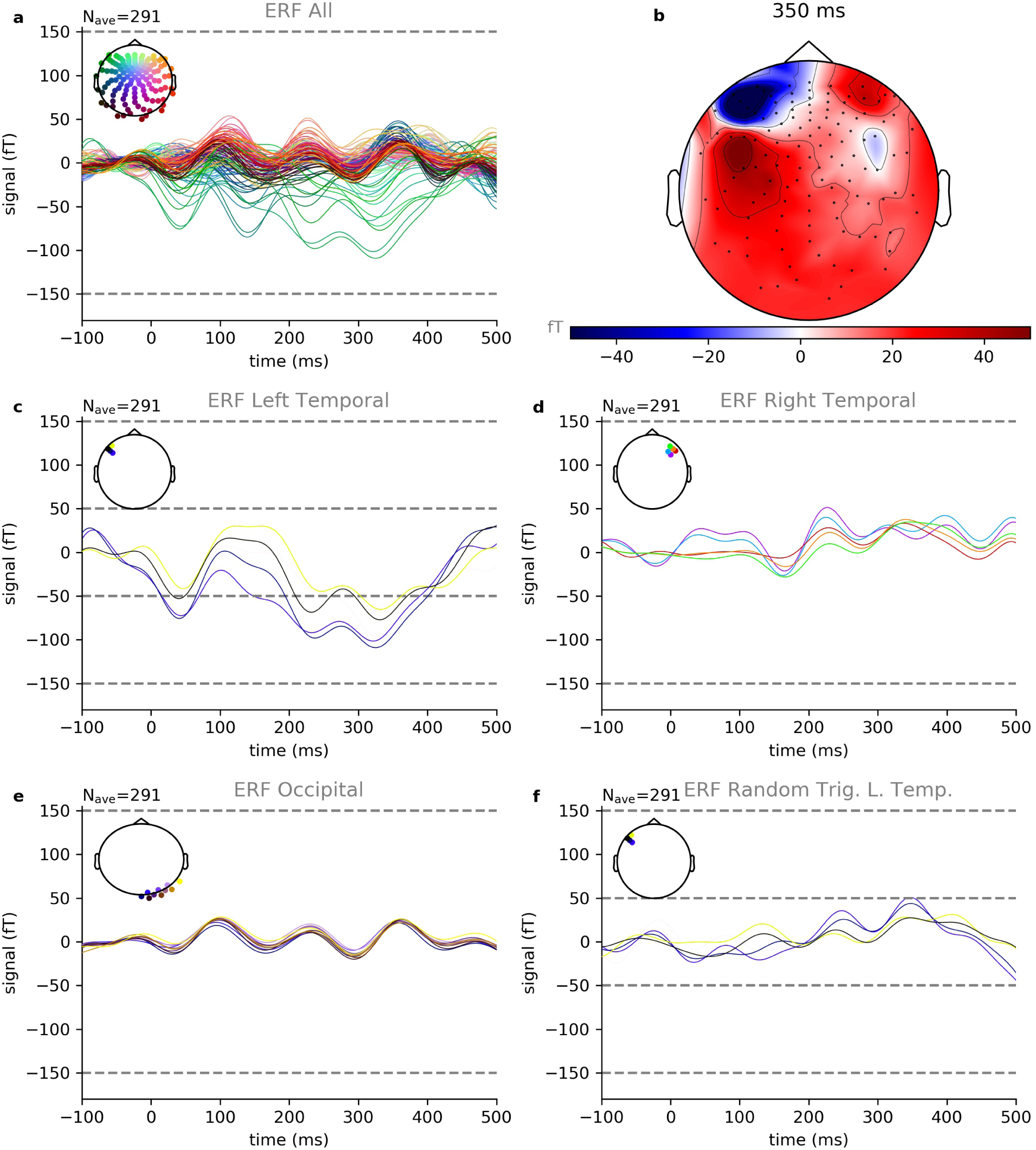
Event related fields for word onset. Shown are exemplary data of book parts number 1-3 of 10 from subject 2 of 15. a: Summary of ERFs of all 248 recording channels averaged over 291 trials. Colors indicate the positions of the recording channels. b: Spatial distribution of ERF amplitudes at 350ms after word onset. c: The largest amplitudes occur in channels located at temporal and frontal areas of the left hemisphere. d: The corresponding channels at the right hemisphere show clearly smaller ERF amplitudes. e: The same is true for occipital channels. f: Same channels as in c, but averaged over randomly chosen triggers instead of word onset triggers. Also in this control condition, the resulting amplitudes are smaller than those for the word onset condition.

**Figure 8:**
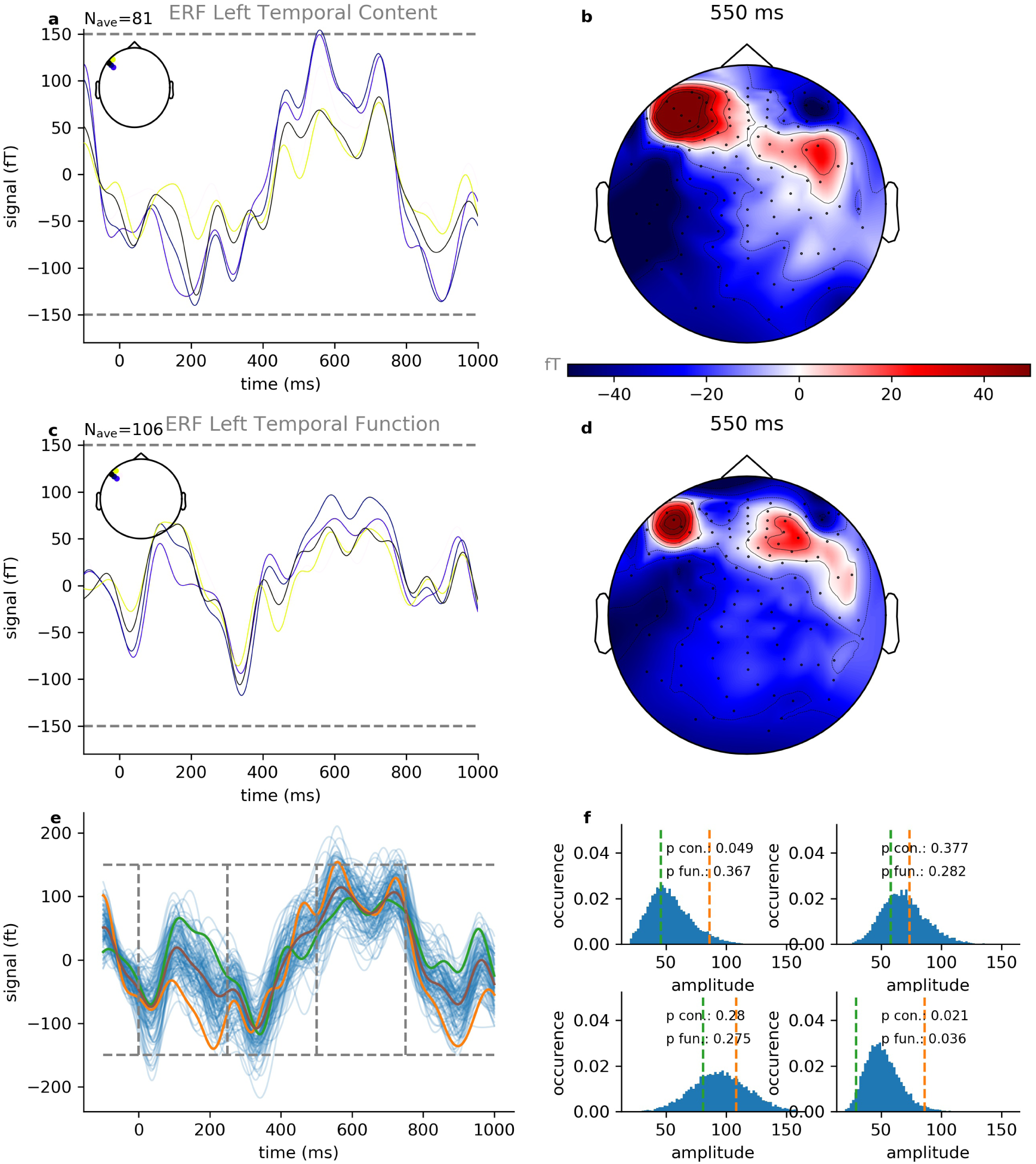
Event related fields for function and content words. Shown are exemplary data of book parts number 1-3 of 10 from subject 2 of 15. a: Averaged ERFs for content words (n=81 trials) with largest amplitudes. b: Spatial distribution of ERF amplitudes at 550ms after word onset for content words. c: Averaged ERFs for function words (n=106 trials) with largest amplitudes. d: Spatial distribution of ERF amplitudes at 550ms after word onset for function words. e: ERF with the largest amplitude for content words (orange) and function words (green), together with ERFs derived from permutation test (blue). f: Distribution of ERF amplitudes derived from permutation test within four subsequent time frames: 0 *ms*–250 *ms* (upper left), 250 *ms*–500 *ms* (upper right), 500 *ms*–750 *ms* (lower left) and 750 *ms*–1000 *ms* (lower right).

Trials with amplitudes higher than 2 · 10^−12^ T would have been rejected, as they were supposed to arise from artifacts. However, none of the trials fit this condition, and hence no trail was rejected.

For this study, we restricted our analyses to sensor space, and did not perform source localization in analogy to other ERF studies [79–81].

### Alignment, segmentation and tagging

Since we have both, the original audio book wave file together with the time tags of word boundaries from forced alignment (Figure 3a), and the corresponding recordings of two analog auxiliary channels of the MEG (Figure 3b), all 248 MEG recording channels could easily be aligned offline with the speech stream (Figure 3c). Subsequently, the continuous multi-channel MEG recordings were segmented using the time tags as boundaries, and labeled with the corresponding types, in our case individual words (Figure 4). Note that, in principle, the process of segmentation can also be performed at different levels of granularity. For instance, using the time tags of phone boundaries would result in a more fine-grained segmentation, whereas grouping several words together to n-grams with appropriate labels to larger linguistic units (i.e. collocations, phrases, clauses, sentences) would result in a more coarse-grained segmentation.

For the analysis of function and content words, we additionally applied POS tagging [60–62] using spaCy [63] to assign word classes (e.g., nouns, verbs, adjectives, conjunctions, determiners, prepositions) to the individual words. According to Ortmann et al. [82], spaCy’s accuracy for POS tagging of German texts is 92.5%. This value could be confirmed by two German native speakers who cross-checked a random sample of sentences, that have been POS tagged using spaCy. However, the most frequent errors observed in spaCy are confusions of nouns and proper names, adverbs and adverbial adjectives, and of different verb forms [82]. Since all these word classes belong to the domain of content words, these confusions are irrelevant for the classification in function and content words analyzed in this study. So that the accuracy for this distinction is expected to be much higher.

### Event related fields

In order to provide the proof-of-principle of our approach, we analysed event related fields (ERF) evoked by word onsets (Figure 7). Since the continuous MEG signals of all 248 channels are already segmented according to word boundaries, we can compute ERFs of word onsets for each channel by simply averaging the pre-processed signals over the word tokens in our database. Here, we included only those words that follow a short pause, instead of using all words occurring in the data set.

Thus, there is a short period of silence, ranging from approximately 50 ms to 1.5 seconds, before the actual word onsets which improves signal quality, yet with the drawback that only a fraction of all tokens can be used. However, there were still 291 remaining events, baseline corrected, within the first three parts of the audio book, corresponding to approx. 12 min of continuous speech, that fit this condition.

In addition, we also analysed ERFs evoked by prototypical content words (nouns, verbs, adjectives) and compared them with ERFs evoked by function words (determiners, prepositions, conjunctions). Again, we included only those words that follow a short interval of silence, instead of using all words occurring in the data set. Within the first three parts of the audio book, this resulted in 81 remaining events for the content word condition and 106 events for the function word condition.^2^

### Permutation test

We performed intra-individual permutation tests [83] to estimate the p-value for the ERF comparison between content and function words. Thus, the ERF was cut into four subsequent time frames, each with a duration of 250 ms, and the root-mean-square amplitude (*RMS*) was calculated (Figure 8e): 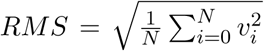, with the signal values within a 250 ms interval *v*_*i*_, the total number of values within a 250 ms interval *N* = *f*_*s*_ · 250 ms, and the sampling rate *f*_*s*_ = 1000 Hz.

10,000 different random permutations of content word and function word labels were generated. For each of these samples the four RMS amplitudes for the different time frames were calculated based on the baseline corrected^3^ single trials, resulting in a distribution of amplitudes for each time frame (Figure 8f). The amplitude values corresponding to the true labeling are compared with amplitudes derived from random permutations in order to estimate the statistical significance, i.e. the p-values for content and function words *p*_*con*_ and *p*_*fun*_, respectively, in Figure 8f.

### Normalized power spectra

Using Fourier transformation, we also analyzed the averaged normalized power spectra (alpha, beta and gamma frequency range) for words in contrast to pauses, and for function and content words. The frequency bands were defined as follows: *α* : 8 − 12 Hz, *β* : 12 − 30 Hz, *γ* : 30 − 45 Hz. The epoch length for this analysis was 400 ms for both, short periods of silence and word onsets. Furthermore, we projected the resulting values to the corresponding spatial position of the sensors. This was done by the usage of the plot psd topomap function of the *MNE* library with Python interface [75, 76].

## Results

The general idea of our approach was to perform MEG measurements of participants listening to an audio book. By synchronizing the continuous speech stream with the ongoing multi-channel neuronal activity, and subsequently automatically segmenting the data streams according to word boundaries derived from forced alignment, we generated a database of annotated speech evoked neuronal activity. This corpus may then be analyzed offline by applying the full range of methods from statistics, natural language processing, and computational corpus linguistics. In order to demonstrate the feasibility of our approach, we restricted our analyses to sensor space and did not perform any kind of source localization (cf. Methods). More specifically, we calculate averaged ERFs for word onsets, and normalized power spectra for onsets of both, words and short pauses. In addition, we compare averaged ERFs for content and function words, and the corresponding normalized power spectra and discuss the results in the light of existing studies.

### Distribution of word classes

We analyzed the distributions of word classes and word class combinations in the audio book and compared them with five different German corpora (German mixed 10k, 30k, and 100k; German news 30k; German Wikipedia 30k) taken from the *Leipzig Corpora Collection* [59], and in addition with a number of other German novels. A sample of the resulting distribtuions is provided in Figure 5. It turns out that *Gut gegen Nordwind* seems to have a very typical word class distribution (Figure 5a,b), especially in comparison to other German novels (Figure 5c-h). In contrast, in the German mixed corpus, there seem to be an under representation of pronouns (Figure 5i,j) compared to all analyzed novels. Using multi-dimensional scaling (MDS), we visualize the mutual (dis-)similarities between all word class distributions (Figure 6a), and distributions of word class combinations (Figure 6b). We find that *Gut gegen Nordwind* is closer, i.e. more similar, to the German novels than to the German corpora. The five corpora seem to cluster apart from the novels. In particular, the distributions of the German mixed corpora of three different sizes (10k, 30k, 100k words) are almost indistinguishable, and hence the corresponding MDS projections are overlapping. Furthermore, it remarkably turns out, that different novels from the same author are closer, i.e. more similar in terms of word class and word class combination distributions, than novels from different authors.

### Event related fields of word onsets

For a first proof of concept and to determine clear neurophysiological brain responses from continuous speech, we analyzed event related fields (ERFs) for word onsets (irrespective of their word classes) from different topographical sides.

Figure 7a shows one example of the resulting ERFs averaged over the aforementioned 291 events corresponding to word onsets for one participant (subject 2 of 15) and parts number 1 to 3 of the audio book and Figure 7b shows a projection of the spatial distribution of the ERF amplitudes at 350 ms after word onset. The largest amplitudes occur in channels located at temporal and frontal areas of the left hemisphere known to be associated with language processing [85]. The ERFs of those channels with the largest ERF amplitudes are shown in Figure 7c. Furthermore, we see a clear N400 component for the word onset condition, indicating language associated processing (cf. [20, 86–90]).

In order to exclude random effects, we compare these channels with the corresponding channels located at the right hemisphere – where we expect less activation due to the asymmetric lateralization of speech in the brain – (Figure 7d), and with some occipital channels (Figure 7e). In both cases, the resulting ERF amplitudes are clearly smaller than those of the left temporal and frontal channels (Figure 7c). In addition, we calculate control ERFs for the same channels shown in Figure 7c, but instead of word boundaries we used randomly chosen time tags for segmentation. Also in this control condition, the resulting ERF amplitudes are smaller than those for the word onset condition (Figure 7f). This result, in particular, demonstrates that even though there are no or only relatively short inter stimulus intervals, leading to overlapping effects of late and early responses of subsequent words, there is still enough signal left in the individual trials.

Finally, we evaluated the re-test reliability of our results using three-fold sub-sampling by separately averaging only over events belonging to the same part of the the audio book (Figures S1, S2, S3). Again, the largest ERF amplitudes were found in the same channels as before and all results show very similar patterns to those shown in Figure 7. In addition, we provide exemplary results of two further participants in the Supplements section (Figures S4 and S5).

### Event related fields of content and function words

As a further validity test of the present study, we analyzed and compared the brain responses of different word classes. As an example, the resulting ERFs averaged over the respective events (content words: n=81, function words: n=106) for one participant (subject 2 of 15) and parts number 1 to 3 of the audio book are shown in Figure 8a,c. and a projection of the spatial distribution of the ERF amplitudes at 550 *ms* after word onset is provided in Figure 8b,d. Again, we see a clear N400 component for both conditions, indicating language associated processing (cf. [20, 86–90]).

Furthermore, we found that content words (Figure 8a,b) elicited greater activation than function words (Figure 8c,d), especially in temporal and frontal areas of the left hemisphere. Since content parts of speech have been shown to differ semantically from function parts of speech [91, 92], these findings are in line with previous studies [57].

In addition, we compared the two conditions for the channel yielding the largest ERF amplitude and performed a permutation test [83] independently for four subsequent time frames each with a duration of 250 ms (Figure 8e,f). We found that the averaged ERFs for the two conditions (content and function words, intra-individual) are significantly (*p* < 0.05) different within the first (0 ms– 250 ms) and third (500 ms–750 ms), but not within the second (250 ms–500 ms) and fourth (750 ms– 1000 ms) time frame (Figure 8f). These results are consistent across all subjects (cf. e.g., Figures S6 and S7 for two further subjects), and are in line with previously reported results [93].

### Averaged normalized power spectra

In our analysis of the averaged normalized power spectra we were unable to find significant differences between the conditions of word onset and of silence onset (Figure 9), and neither between content words and function words (Figure 10). See discussion section for possible reasons.

**Figure 9:**
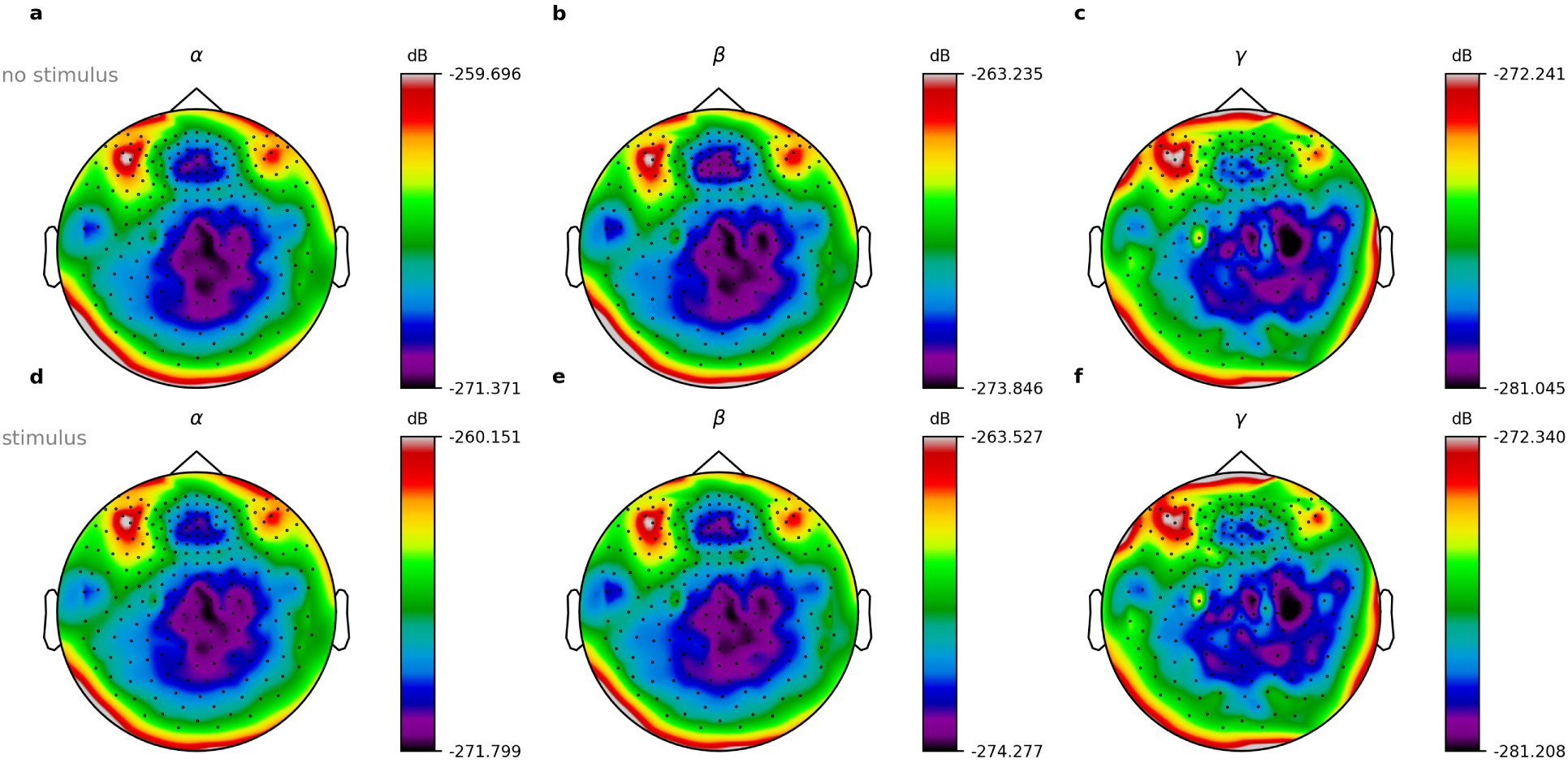
Normalized power spectra for words and silence. Shown are exemplary data of book parts number 1-3 of 10 from subject 2 of 15. a-c: Power spectra for word offset, i.e. silence. d-f: Power spectra for word onsets. a,d: Alpha frequency range. b,e: Beta frequency range. c,f: Gamma frequency range.

**Figure 10:**
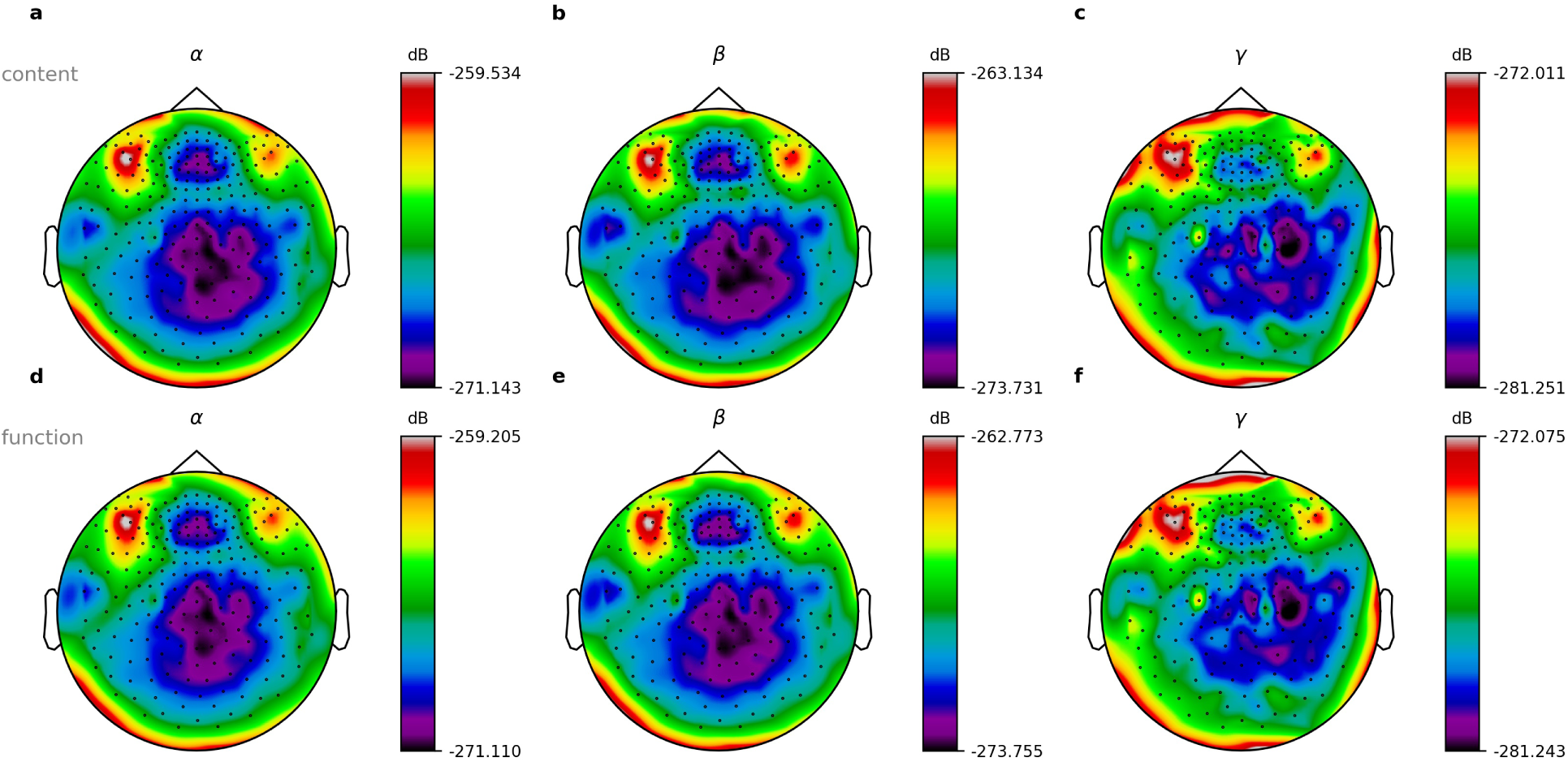
Normalized power spectra for content and function words. Shown are exemplary data of book parts number 1-3 of 10 from subject 2 of 15. a-c: Power spectra for content words. d-f: Power spectra for function words. a,d: Alpha frequency range. b,e: Beta frequency range. c,f: Gamma frequency range.

## Discussion

In this study, we presented an approach where we combine electrophysiological assessment of neuronal activity with computational corpus linguistics, in order to create a corpus as defined in [62] of continuous speech-evoked neuronal activity. We demonstrated that using an audio book as natural speech stimulus, and simultaneously performing MEG measurements led to a relatively large number of analysable events (word onsets: *n* = 291, silence onsets: *n* = 187, content words: *n* = 81, function words: *n* = 106), yet within a relatively short measurement time of 15 *min*. We further provided the proof-of-principle that, in contrast to common study designs, even though our stimulus trials were not presented in isolation, i.e. with appropriate inter-stimulus intervals of a few seconds, averaging over all respective events of a certain condition results in ERFs in left temporal and frontal channels with increased amplitudes compared to those of several control channels (e.g., at right hemisphere or at occipital lobe). The same is true with respect to comparison with control conditions (e.g., random trigger times). These results are well in line with previously published findings [85].

Furthermore, we analyzed ERFs for different categories of words. Although, a frequently investigated and contrasted pair of word classes is that of nouns and verbs [94–99], for the present study, we opted for the distinction between function words, defined as determiners, prepositions and conjunctions, and content words, defined as nouns, verbs and adjectives. These lexical categories are also frequently used in neuroimaging studies on the neurobiology of language [9, 57, 93, 100–102]. In addition, they differ greatly in the semantic domain, and cover more fully the the totality of the words than the categories of nouns and verbs, since nouns and verbs are both included in the content word category. We found a clear N400 component (cf. Figure 8) especially in left hemispheric frontal regions for both function and content words and a positive component from 400-700 ms which is in line with Brennan et al.’s findings [103,104]. Additionally, we found that content words elicit greater activation than function words, especially in temporal and frontal areas of the left hemisphere. Due to their substantial semantic differences [91, 92], this finding is in line with previous studies [57].

With respect to the average normalized power spectra, it was found that presentation of speech stimuli was associated with an increase in broadband gamma and a decrease in alpha over auditory cortex, while alpha power was increased in domain unspecific cortical areas [105–107]. One reason could be that, since we analysed only very short periods of silence, i.e. between two words, our two conditions of word onset and silence onset can be considered basically, at a larger time scale, to be the same condition, i.e. continuous speech stimulation. This may explain why we found no differences in frequency power here. Even though it has been proposed that in human language networks linguistic information of different types is transferred in different oscillatory bands – in particular attention is assumed to correlate with an increase in gamma and a decrease in alpha band power [108] – the role of different spectral bands in mediating cognitive processes is still not fully understood. Therefore it remains unclear, whether these findings extend to content and function words. Whether our approach is too insensitive to see differences here remains to be seen, and further studies should look more closely at this issue.

As mentioned above, in contrast to traditional studies that are limited to testing only a small number of stimuli or word categories, the present approach opens the possibility to explore the neuronal correlates underlying different word meaning information across a large range of semantic categories [3], and syntactic structures [109]. This is because the ongoing natural speech used here contains both, a large number of words from different semantic domains [37] and a large number of sentences at all levels of linguistic complexity [110].

On the other hand, one may argue that stimulation with ongoing natural speech has, compared to traditional approaches, the drawback that there are virtually no inter stimulus intervals between the single words. This, of course, introduces a mixture of effects at different temporal scales, e.g. early responses to the actual word are confounded with late responses of the previous word. However, all these effects may be averaged out, as demonstrated by other studies [4, 18–21] and also by our results.

In a follow-up study, it will have to be validated whether our approach also works for linguistic units of different complexity other than single words. For instance, smaller linguistic units like phonemes and morphemes, but also larger linguistic units like collocations, phrases, clauses, sentences, or even beyond, could be investigated. For instance, we might be able to determine what neural correlates of the different association measures used in research on collocation look like (see [111] for an overview and further references). Furthermore, more abstract linguistic phenomena need to be analyzed, e.g. argument structure constructions [112–114] or valency [115–117]. Finally, our speech-evoked neural data may also be grouped, averaged, and subsequently contrasted according to male and female voice, looking at gender specific differences (see e.g., [118, 119]).

Also, analyses based on source space need to be tested, as well as more sophisticated analyses taking advantage of the multi-dimensionality of the data, such as, for instance, multi-dimensional cluster statistics [120, 121]. In addition, state-of-the-art deep learning approaches may be used as a tool for analyzing brain data, e.g. for creating so-called *embeddings* of the raw data [122]. Moreover, as proposed by Kriegeskorte [123], our neural corpus can serve to test [124] computational models of brain function [125–128], in particular models based on neural networks [129–131] and machine learning architectures [132,133], in order to iteratively increase biological and cognitive fidelity [123].

Due to the corpus-like features of our data, all additional analyses mentioned may be performed on the existing database, and without the need for designing new stimulation paradigms, or carrying out additional measurements.

However, in order to avoid statistical errors due to *HARKing* [54, 55] - defined as generating scientific statements exclusively based on the analysis of huge data sets without previous hypotheses - and to guarantee consistency of the data, it is necessary to apply e.g., re-sampling techniques such as sub-sampling as shown above and described in detail in [134]. Furthermore, the approach presented here allows us to apply the well-established machine learning practice of data set splitting, i.e. to split the dataset into multiple parts before the beginning of the evaluation, where the one part is used for generating new hypotheses, and another part for subsequently testing these hypotheses (or split again into training and testing data). However, since we recorded a whole story, possible order effects should be taken into account for dataset splitting. Hence, instead of splitting the data set according to the chronological order, e.g. using the first parts of the audio book as training, and the subsequent parts as test dataset, it should better be split randomly.

To conclude, there are two major reasons why we think the study of the neurobiology of language can benefit tremendously from the introduction of corpus-linguistic methodology.

The first is that we can base our research on naturally occurring language, which should make them more ecologically valid than the more artificial stimuli used in carefully balanced and controlled experiments. Of course, even though audio books are frequently used in similar studies [4, 18–21], one may also discuss whether audio books actually can be considered natural speech. One could argue that the fact that highly trained professional speakers and actors are usually employed to read audio books, who may use specific intonational patterns to paint a more vivid image of the situation, may lead to unnaturalness and thus possibly to unusual arousal patterns in the hearer. However, this argument is flawed. People spend large portions of their days listening to language produced by such professional speakers for radio, television news and drama, online videos, and podcasts. While probably not predominant for most people, it corresponds to a perfectly normal, everyday type of language experience. Even if we expect deviations from spoken interaction in such stimuli, we could even exploit this to study brain responses to creative language use (see [135], [136] and the sources cited there for linguistic studies of creativity). Of course, further studies using recordings of everyday dialogues between untrained subjects, e.g. describing what they have done during the day, should be designed to obtain a more comprehensive picture and more robust results, because, as Kriegeskorte and Douglas pointed out that *“as we engage all aspects of the human mind, our tasks will need to simulate natural environments”* [123]. Still, purely receptive task such as the one used in this study is one type of natural environment, and one that can be studied without too much interference compared to, say, spontaneous interaction.

The second reason is the fact that measurements can be re-used if they form part of a large corpus of neuroimaging results. Let us look at a few numbers: In the present study, we stimulated 15 participants with 40 minutes of audio each. Test time spent in the MEG was 60 minutes due to the questions and pauses mentioned above. With 30 minutes of preparation, we used the MEG lab for a total of 22.5 hours during experimentation. In that period of time, we gathered measurements for roughly 6,000 words perceived by 15 participants, totalling 90,000 sets of brain responses to words. These correspond to roughly 35 GB of measurements (4 bytes per value, 1,000 per second, 248 channels, 40 minutes per participant, 15 participants). For this study, we only looked at a tiny fraction of the data (words preceded by a short pause of at least 50 ms in the first 12 minutes) and already managed to confirm certain patterns found by previous studies with a strict experimental design. If we assume that pauses are equally distributed across the corpus, we can expect to find roughly 1,000 such events, with 15 participants for each, i.e. 15,000 data points alone for words preceded by silence. Having these plus all the other words in their immediate linguistics contexts without pauses opens many more avenues for interesting research question at no added laboratory costs. Once we start looking at all words, we expect that the noise introduced through not being able to control for a variety of factors will be counterbalanced by the sheer size of data sets constructed using the methodology presented.

By that, we agree with the view of Hamilton and Huth that *“natural stimuli offer many advantages over simplified, controlled stimuli for studying how language is processed by the brain”*, and that *“the downsides of using natural language stimuli can be mitigated using modern statistical and computational techniques”* [137].

## Acknowledgments

This work was funded by the Deutsche Forschungsgemeinschaft (DFG, German Research Foundation): grant KR5148/2-1 to PK – project number 436456810, the Interdisciplinary Center for Clinical Research (IZKF) at the University Hospital of the University Erlangen-Nuremberg (grant ELAN-17-12-27-1-Schilling to AS), and the Emergent Talents Initiative (ETI) of the University Erlangen-Nuremberg (grant 2019/2-Phil-01 to PK).

We are grateful to the publishers *Deuticke Verlag* and *Hörbuch Hamburg* for the permission to use the novel and corresponding audio book *Gut gegen Nordwind* by Daniel Glattauer for the present and future studies.

We thank Martin Kaltenhäuser for technical assistance, and Stefan Rampp for useful discussion.

Finally, we wish to thank the anonymous reviewers for their remarks and advice which significantly increased the value of our work.

## Additional Information

### Data availability statement

Data will be made available to other researches on reasonable request.

### Author contributions

PK and AS designed the study. AS, PK, AZ and VK prepared the stimulation and processed the audio book. AS, PK, VK and MH performed the experiments. AS and PK analyzed the data. AS, PK, AM, AZ and KS analyzed the text of the novel. AS, PK, RT, MRHS, AM and PU discussed the results. PK, AS, RT, MRHS and PU wrote the manuscript.

## Supplements

**Figure S1:**
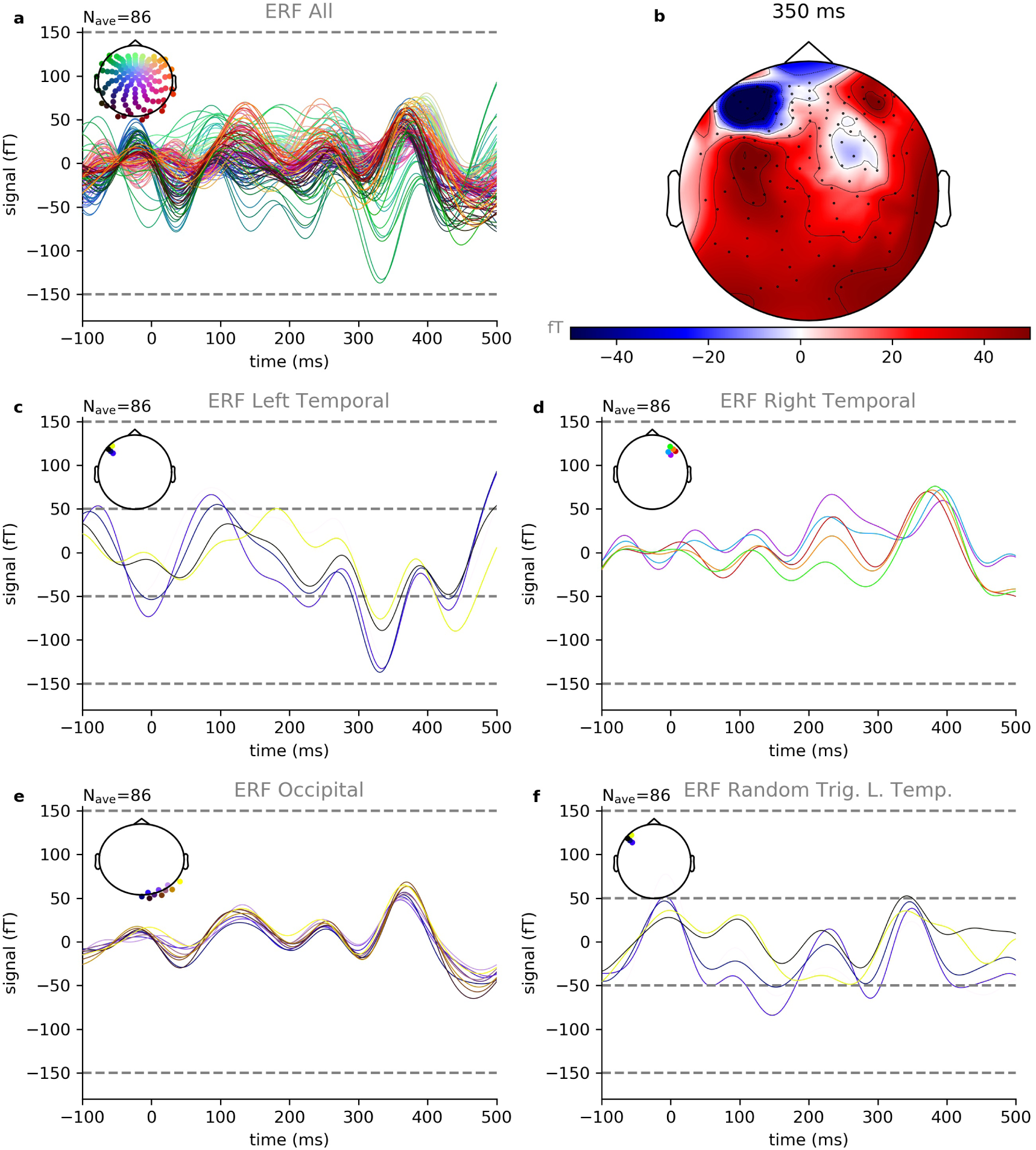
Event related fields of word onsets. Sub-sample 1 of 3. Shown are exemplary data of book part number 1 of 10 from subject 2 of 15. a: Summary of ERFs of all 248 recording channels averaged over 86 trials. Colors indicate the positions of the recording channels. b: Spatial distribution of ERF amplitudes at 350ms after word onset. c: The largest amplitudes occur in channels located at temporal and frontal areas of the left hemisphere. d: The corresponding channels at the right hemisphere show clearly smaller ERF amplitudes. e: The same is true for occipital channels. f: Same channels as in c, but averaged over randomly chosen triggers instead of word onset triggers. Also in this control condition, the resulting amplitudes are smaller than those for the word onset condition.

**Figure S2:**
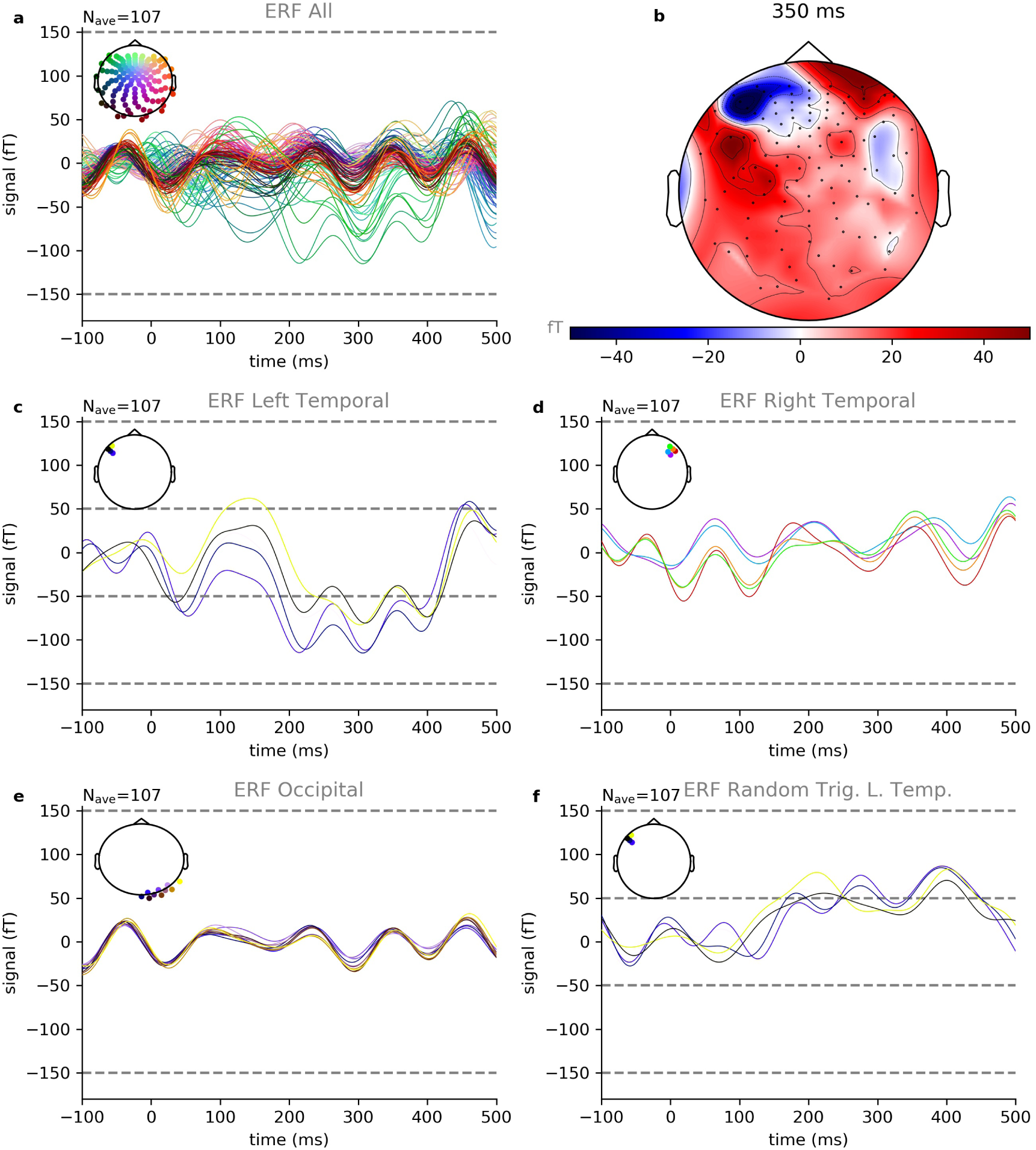
Event related fields of word onsets. Sub-sample 2 of 3. Shown are exemplary data of book part number 2 of 10 from subject 2 of 15. a: Summary of ERFs of all 248 recording channels averaged over 107 trials. Colors indicate the positions of the recording channels. b: Spatial distribution of ERF amplitudes at 350ms after word onset. c: The largest amplitudes occur in channels located at temporal and frontal areas of the left hemisphere. d: The corresponding channels at the right hemisphere show clearly smaller ERF amplitudes. e: The same is true for occipital channels. f: Same channels as in c, but averaged over randomly chosen triggers instead of word onset triggers. Also in this control condition, the resulting amplitudes are smaller than those for the word onset condition.

**Figure S3:**
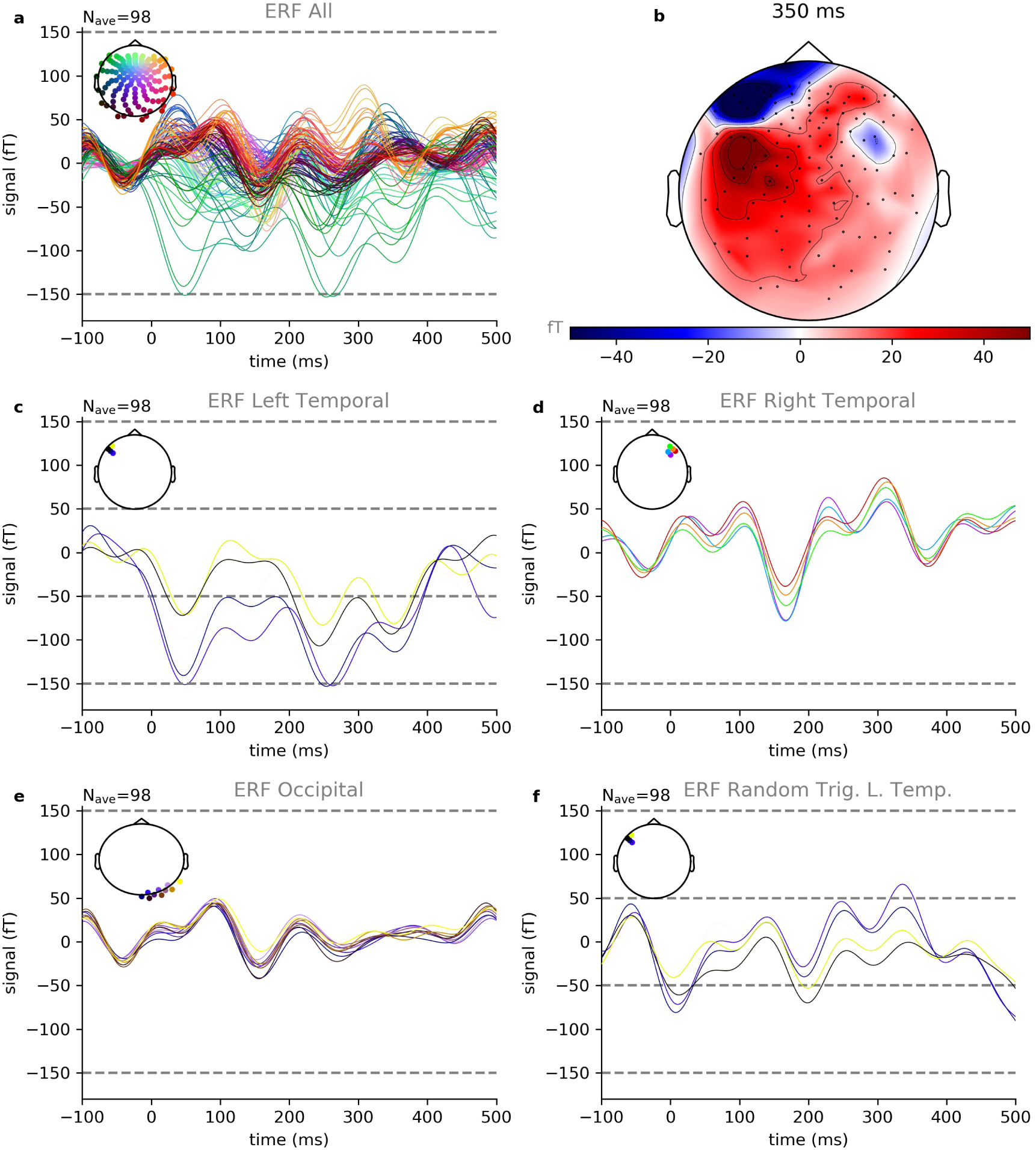
Event related fields of word onsets. Sub-sample 3 of 3. Shown are exemplary data of book part number 3 of 10 from subject 2 of 15. a: Summary of ERFs of all 248 recording channels averaged over 98 trials. Colors indicate the positions of the recording channels. b: Spatial distribution of ERF amplitudes at 350ms after word onset. c: The largest amplitudes occur in channels located at temporal and frontal areas of the left hemisphere. d: The corresponding channels at the right hemisphere show clearly smaller ERF amplitudes. e: The same is true for occipital channels. f: Same channels as in c, but averaged over randomly chosen triggers instead of word onset triggers. Also in this control condition, the resulting amplitudes are smaller than those for the word onset condition.

**Figure S4:**
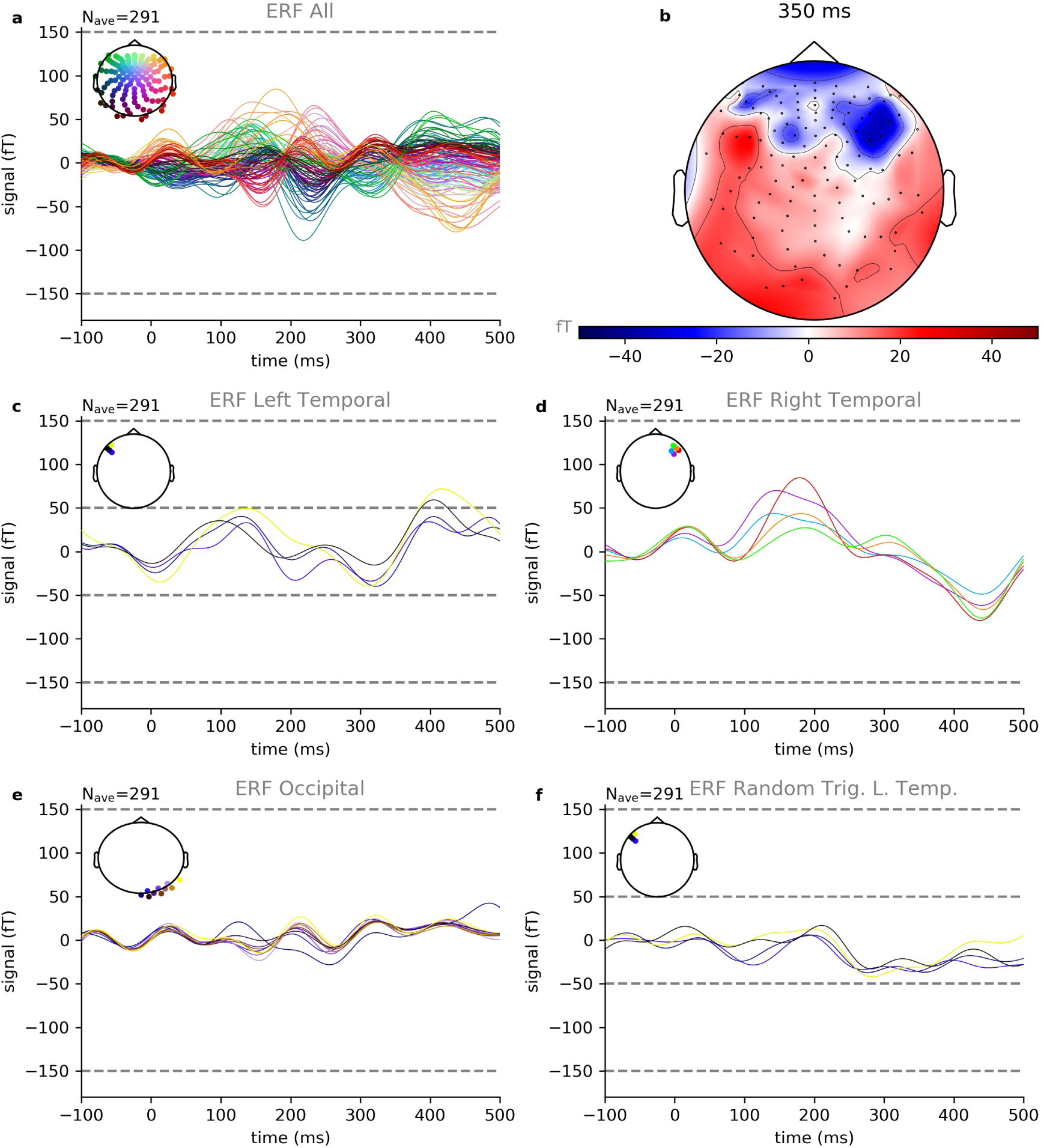
Event related fields for word onset. Shown are exemplary data of book parts number 1-3 of 10 from subject 5 of 15. a: Summary of ERFs of all 248 recording channels averaged over 291 trials. Colors indicate the positions of the recording channels. b: Spatial distribution of ERF amplitudes at 350ms after word onset. c: The largest amplitudes occur in channels located at temporal and frontal areas of the left hemisphere. d: The corresponding channels at the right hemisphere show clearly smaller ERF amplitudes. e: The same is true for occipital channels. f: Same channels as in c, but averaged over randomly chosen triggers instead of word onset triggers. Also in this control condition, the resulting amplitudes are smaller than those for the word onset condition.

**Figure S5:**
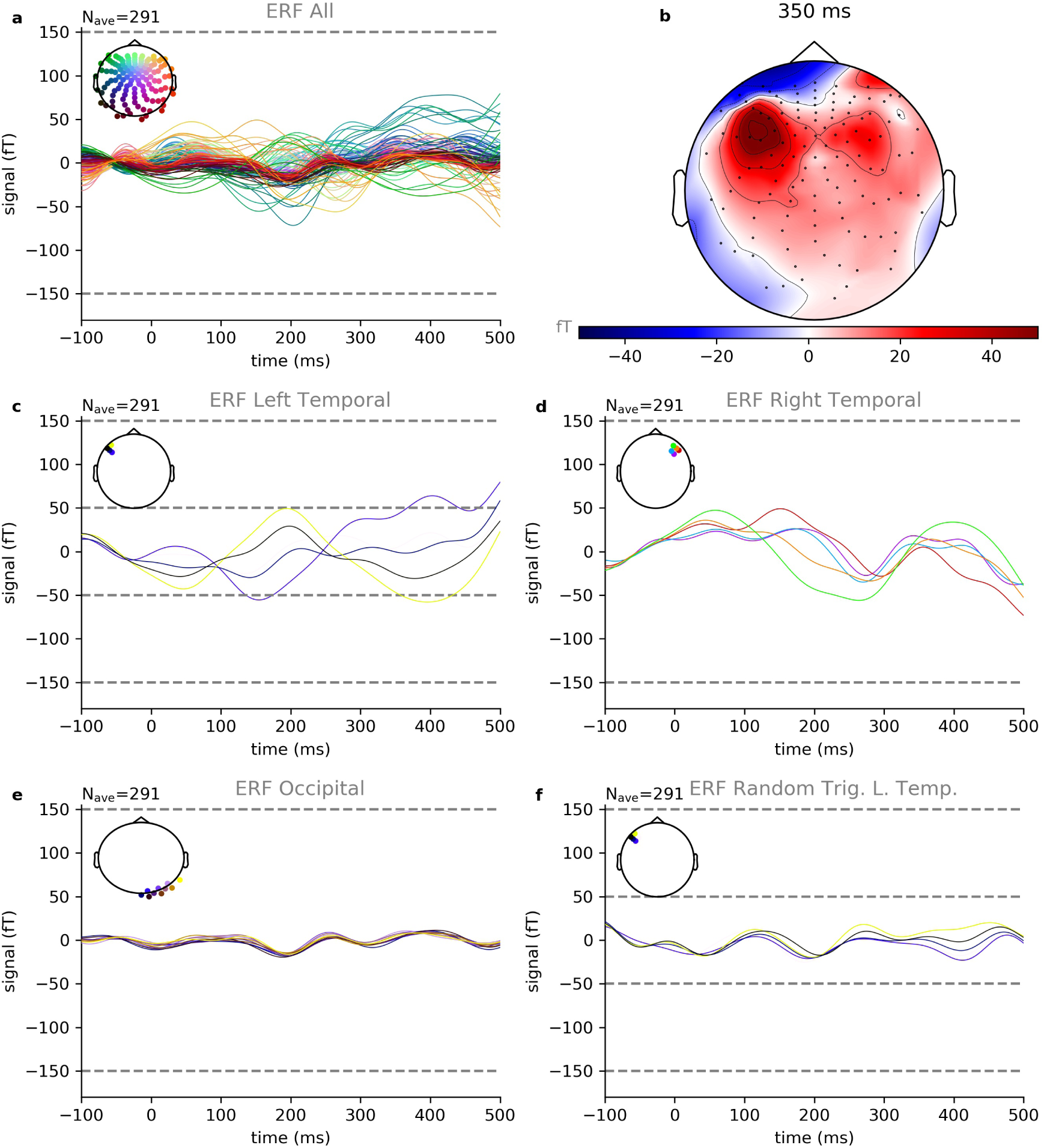
Event related fields for word onset. Shown are exemplary data of book parts number 1-3 of 10 from subject 8 of 15. a: Summary of ERFs of all 248 recording channels averaged over 291 trials. Colors indicate the positions of the recording channels. b: Spatial distribution of ERF amplitudes at 350ms after word onset. c: The largest amplitudes occur in channels located at temporal and frontal areas of the left hemisphere. d: The corresponding channels at the right hemisphere show clearly smaller ERF amplitudes. e: The same is true for occipital channels. f: Same channels as in c, but averaged over randomly chosen triggers instead of word onset triggers. Also in this control condition, the resulting amplitudes are smaller than those for the word onset condition.

**Figure S6:**
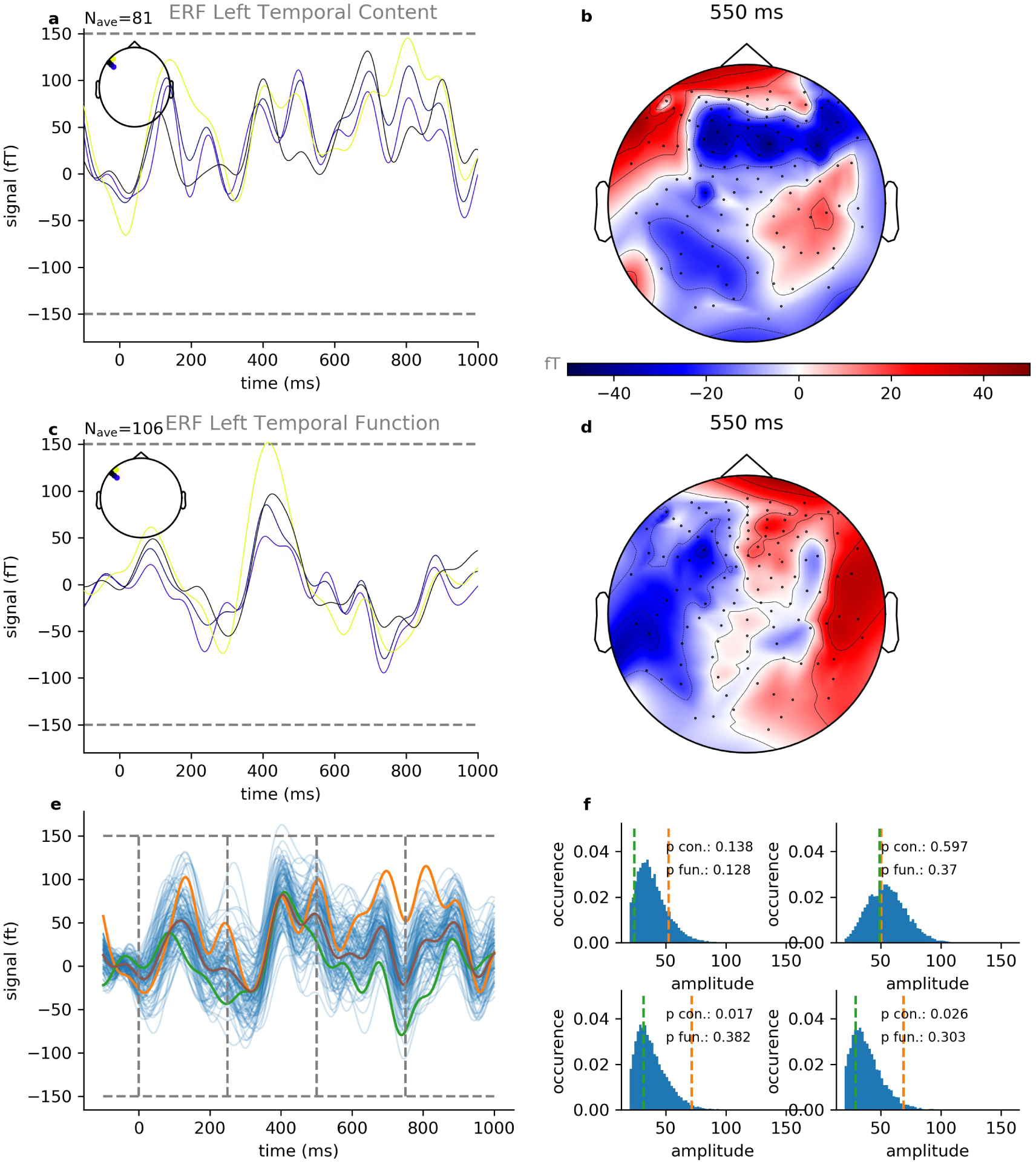
Event related fields for function and content words. Shown are exemplary data of book parts number 1-3 of 10 from subject 5 of 15. a: Averaged ERFs for content words (n=81 trials) with largest amplitudes. b: Spatial distribution of ERF amplitudes at 550ms after word onset for content words. c: Averaged ERFs for function words (n=106 trials) with largest amplitudes. d: Spatial distribution of ERF amplitudes at 550ms after word onset for function words. e: ERF with the largest amplitude for content words (orange) and function words (green), together with ERFs derived from permutation test (blue). f: Distribution of ERF amplitudes derived from permutation test within four subsequent time frames: 0 *ms*–250 *ms* (upper left), 250 *ms*–500 *ms* (upper right), 500 *ms*–750 *ms* (lower left) and 750 *ms*–1000 *ms* (lower right).

**Figure S7:**
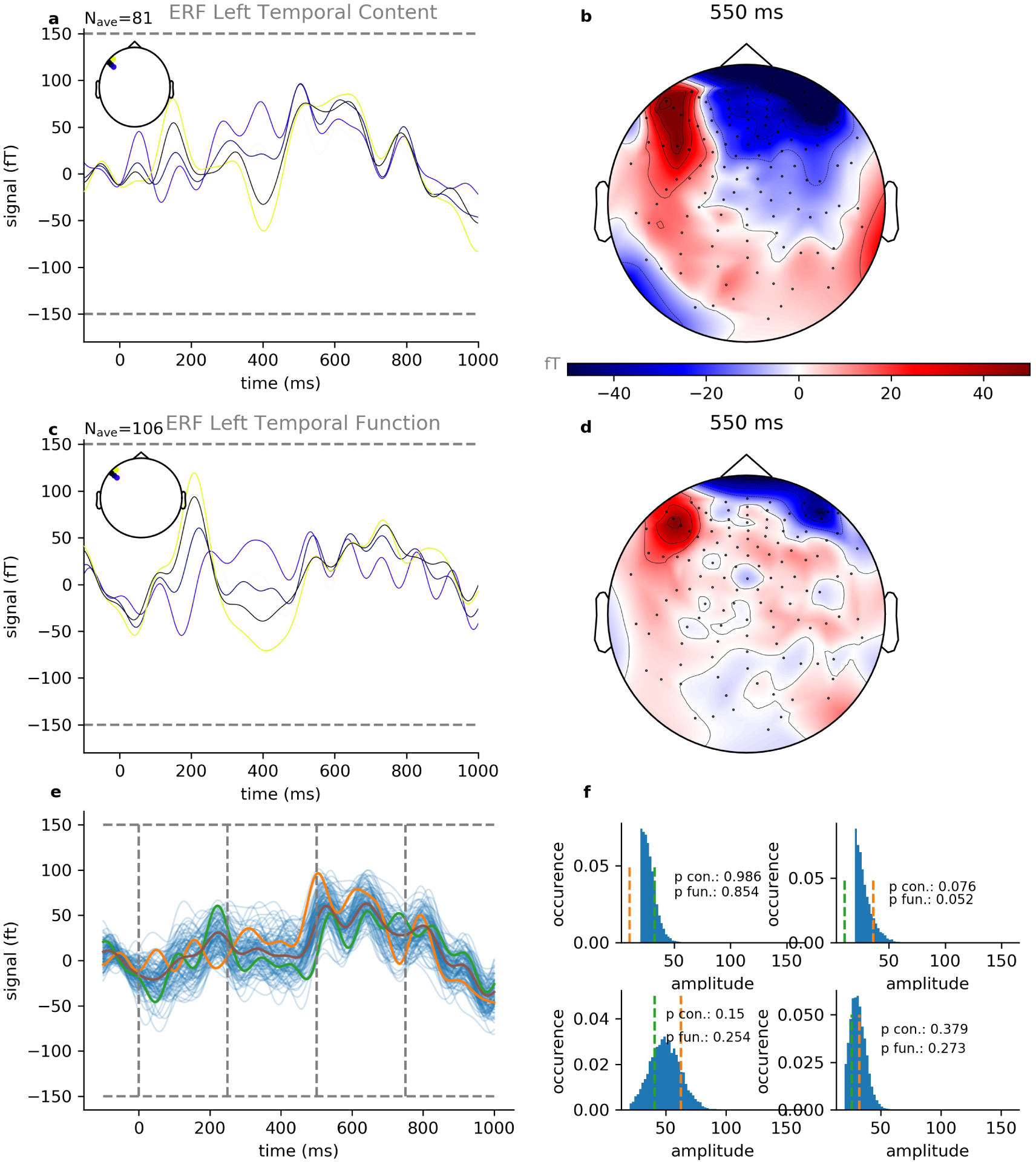
Event related fields for function and content words. Shown are exemplary data of book parts number 1-3 of 10 from subject 8 of 15. a: Averaged ERFs for content words (n=81 trials) with largest amplitudes. b: Spatial distribution of ERF amplitudes at 550ms after word onset for content words. c: Averaged ERFs for function words (n=106 trials) with largest amplitudes. d: Spatial distribution of ERF amplitudes at 550ms after word onset for function words. e: ERF with the largest amplitude for content words (orange) and function words (green), together with ERFs derived from permutation test (blue). f: Distribution of ERF amplitudes derived from permutation test within four subsequent time frames: 0 *ms*–250 *ms* (upper left), 250 *ms*–500 *ms* (upper right), 500 *ms*–750 *ms* (lower left) and 750 *ms*–1000 *ms* (lower right).

**Figure S8:**
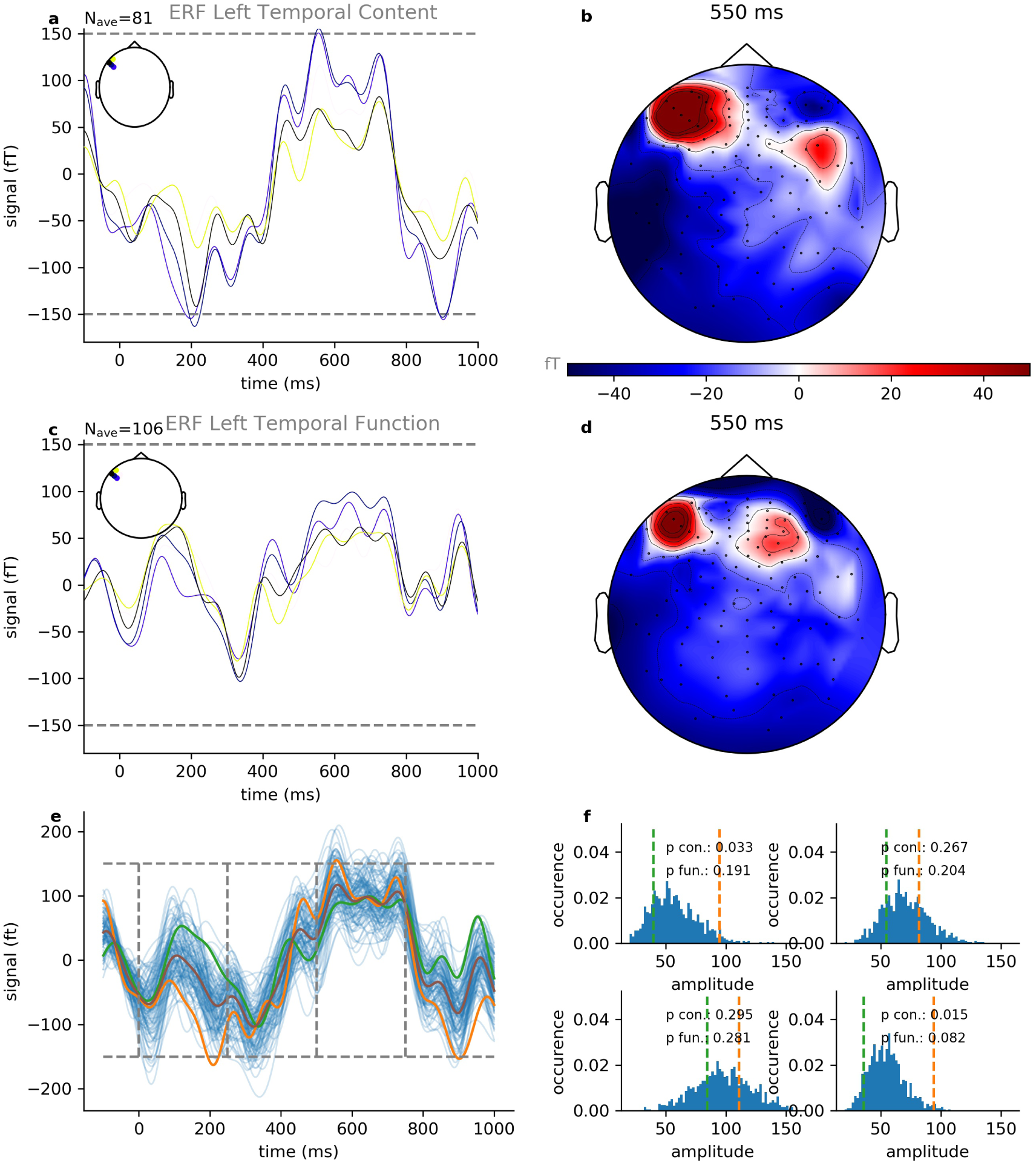
Event related fields for function and content words. Shown are the same analyses and participant as in Figure 8. However, for these analyses we did not delete the first two independent components. As the results are qualitatively equal, this does not affect the difference of neural responses to content and function words.

## Footnotes

1 We follow the categorization of these studies and thus only include the content and function word classes listed above in our study.

2 The numbers do not add up to the total of 291 words because only the word classes listed in the introduction were included in the analysis of content and function words.

3 Following the suggestions published by Alday, instead of traditional baseline correction, we performed strong high-pass filtering with a cutoff frequency of 0.1 Hz, since traditional baseline correction eventually reduces signal-to-noise ratio, and seems therefore to be statistically unnecessary or even undesirable [84].

